# Clonal reconstruction from time course genomic sequencing data

**DOI:** 10.1101/832063

**Authors:** Wazim Mohammed Ismail, Haixu Tang

**Affiliations:** School of Informatics, Computing and Engineering, Indiana University, Bloomington, IN, USA

**Author notes:** Equal contributor.

**Keywords:** clonal reconstruction, time course, maximum likelihood, long-term evolution experiment

## Abstract

**Background:** Bacterial cells during many replication cycles accumulate spontaneous mutations, which result in the birth of novel clones. As a result of this *clonal expansion*, an evolving bacterial population has different clonal composition over time, as revealed in the long-term evolution experiments (LTEEs). Accurately inferring the *haplotypes* of novel clones as well as the clonal frequencies and the clonal evolutionary history in a bacterial population is useful for the characterization of the evolutionary pressure on multiple correlated mutations instead of that on individual mutations.

**Results:** In this paper, we study the computational problem of reconstructing the haplotypes of bacterial clones from the *variant allele frequencies* observed from an evolving bacterial population at multiple time points. We formalize the problem using a maximum likelihood function, which is defined under the assumption that mutations occur spontaneously, and thus the likelihood of a mutation occurring in a specific clone is proportional to the frequency of the clone in the population when the mutation occurs. We develop a series of heuristic algorithms to address the maximum likelihood inference, and show through simulation experiments that the algorithms are fast and achieve near optimal accuracy that is practically plausible under the maximum likelihood framework. We also validate our method using experimental data obtained from a recent study on long-term evolution of Escherichia coli.

**Conclusion:** We developed efficient algorithms to reconstruct the clonal evolution history from time course genomic sequencing data. Our algorithm can also incorporate clonal sequencing data to improve the reconstruction results when they are available. Based on the evaluation on both simulated and experimental sequencing data, our algorithms can achieve satisfactory results on the genome sequencing data from long-term evolution experiments.

**Availability:** The program (ClonalTREE) is available as open-source software on GitHub at https://github.com/COL-IU/ClonalTREE

## Background

Long-term evolution experiment (LTEE) has long been adopted to study how genetic variations are generated and maintained in a period of time and how novel variations are associated with the adaptation of the species to novel environmental conditions [1]. Due to their high genetic diversity and rapid evolution, unicellular microbes, predominantly *E. coli*, are used in LTEEs [2, 3, 4], although LTEE was also conducted on multi-cellular model animals such as Drosophila [5]. The *E. coli* long-term evolution experiment conducted by Lenski and colleagues is the longest on-going LTEE, in which twelve initially identical *E. coli* strains (i.e., the *founder* clones) were grown in parallel, each under a daily serial passage for 30 years [6, 7, 3]. A variety of phenotypic changes were observed in the bacterial population during the experiment, including increased fitness to specific growth conditions [8] and elevated mutation rates [9].

In recent years, LTEE were combined with *metagenome sequencing* (i.e., sequencing the whole genomes in the population, also referred to as the Pool-Seq, or the sequencing of pooled individual genomes) to characterize genetic variations introduced during the course of experiment, and the allele frequencies of these variations in a population [3, 10]. Some of these novel variations were revealed to be associated with observed phenotypic changes, e.g., the defective mutations in the DNA repair pathways causing elevated mutation rates [9], and the novel genetic traits selected for citrate use [10]. Furthermore, population-wide metagenome sequencing can be conducted on the evolving population at multiple time points to monitor the dynamic changes of genetic variations in complex and heterogeneous growth environments. The main objective of these studies is to identify clones adapted to specific environmental niche over the time course. However, due to the nature of metagenome sequencing, it is not straightforward to determine the *haplotypes* of the clones arising in the experiment. Instead, selections are often detected on the novel variations by applying statistical tests [11, 12] to the time series allele frequencies derived from the sequencing data. Because a novel variation, e.g., a single nucleotide variation (SNV), may be shared by multiple clones in the population (i.e. subsequent mutations may occur in a clone already containing mutations instead of the founder clone), the tests on variation may be less sensitive than the tests directly on the frequency profiles of haplotypes, and thus may miss the selection on some clones, especially when the population is dominated by a few clones containing many variations.

To address this issue, in a recent study, metagenome sequencing coupled with clonal sequencing was adopted for the study of populations of wild-type (WT) and repair-deficient *E. coli* evolving over three years [4]. To characterize the haplotypes in the populations, whole genome sequencing was carried out on randomly selected clones at the end of the experiments. In addition, the haplotype frequencies of the major clones were derived from the metagneome sequencing data. The dynamic changes of these major clones during the course of experiment showed a clear picture of the subpopulation structure (e.g., using a Muller plot; see Fig 1C), in which the major clones evolved with different genotypes associated with nutrition metabolism. Despite the demonstrated success here, the clonal sequencing has two disadvantages in practice. First, because the sequenced clones are randomly selected, minor clones with low abundances in the population may not be characterized (while major clones are sequenced repetitively), and thus their frequency profiles during the time course will not be considered in the subsequent analyses. More importantly, the clones are usually chosen at the end of the experiment; as a result, the clones with high abundance in the middle but becoming less abundant towards the end of the experiment are less likely to be characterized, which will not only miss some clones under selection during the time course, but also miscalculate the allele frequencies of characterized clones in the middle of time course. Therefore, unless the clonal sequencing covers a large number of clones (that may contain many duplicated clones) compared to the complexity of the population, it is desirable to develop computational methods to reconstruct the haplotypes of clones from time series metagenome sequencing data.

**Figure 1.**
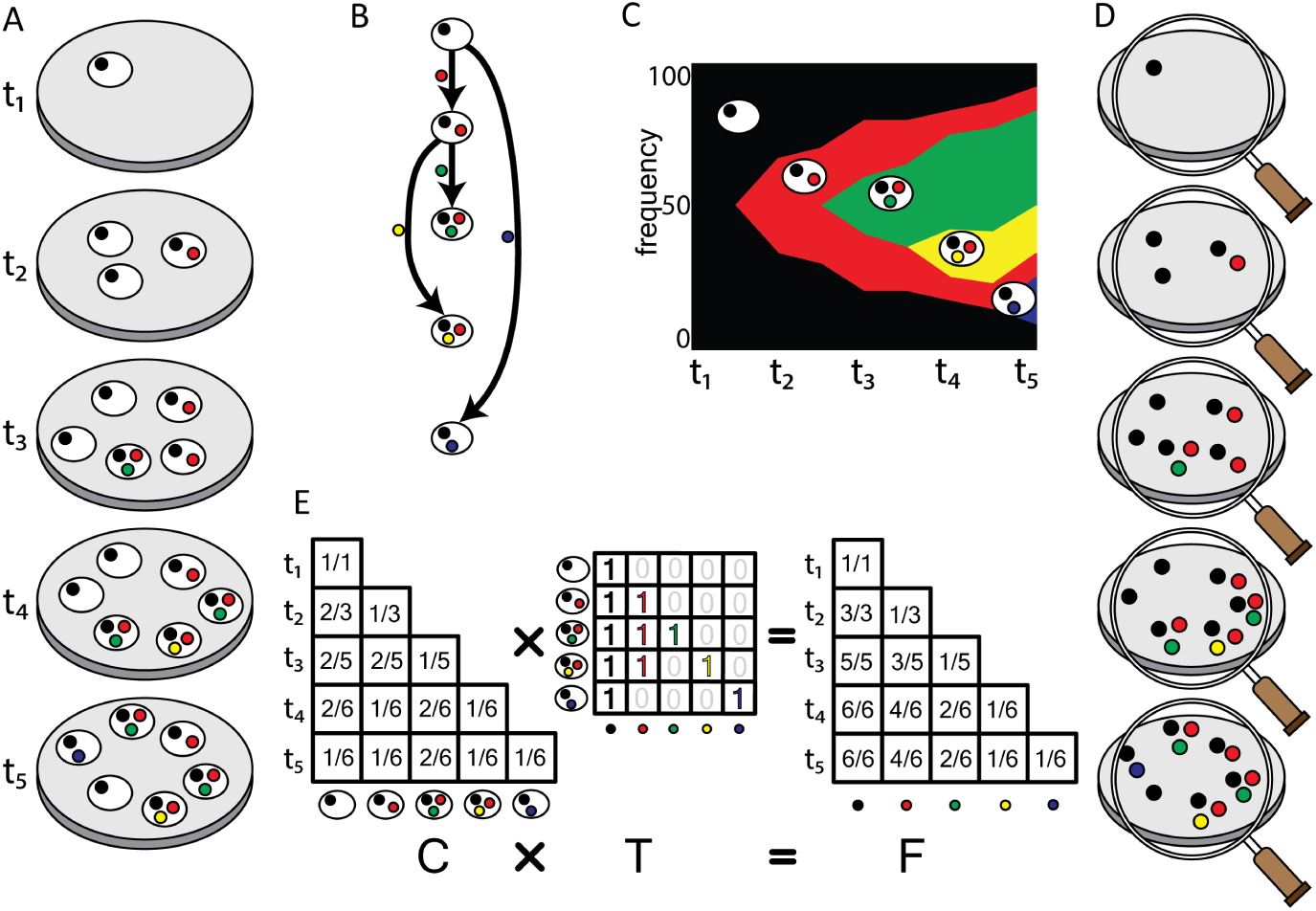
A schematic illustration of the clonal structure in an evolving bacterial population and the time course clonal reconstruction problem. (a) Starting from a single founder clone (shown in black) at time *t*_1_, four mutations (shown in red, green, yellow and blue, respectively) occur at time point *t*_2_ to *t*_5_, respectively, resulting in four novel clones (denoted by their unique variants). (b) The clonal tree represents the evolutionary history of these clones, in which each node represents a clone including the founder clone as the root, and each edge represents the mutations that occur at specific time points. (c) The Muller plot shows the evolutionary dynamics with the novel clones along with their frequencies at each time point. (d) Metagenome sequencing conducted at different time point, from which the variant allele frequencies (VAF) matrix can be derived. (e) The VAF matrix can be viewed as the product of the clonal tree (*T*) and the clone frequencies (*C*), similar to the formulation in cancer genomics [16]. The goal of this work is to reconstruct the clonal tree (*T*) and the clone frequencies (*C*) from the observed VAF matrix.

Interestingly, the clonal reconstruction has been extensively studied in the field of cancer genomics for tracking the evolution of cancer cells by bulk tumor genome sequencing [13, 14], in an attempt to characterize the intra-tumor heterogeneity (i.e., clonal tree and composition) and in the mean time to identify the clones carrying *driver mutations* that occur in the early stage of cancer and drive the cancer progression [15]. Computationally, the *clonal reconstruction* (also referred to as the *clonality inference*) takes as input the allele frequencies of a set of genetic variants in multiple samples (e.g., dissected from the same tumor tissue), and aims to reconstruct a set of clones, each carrying a subset of the variants, and simultaneously infer the fraction of these clones in each sample [16]. Many algorithms addressed the clonal reconstruction problem [17, 18, 19, 20, 16, 21] by inferring the evolutionary history of reconstructed clones and the generation of variants (assuming that each variant is generated only once, i.e., the *infinite sites assumption* [22]), from which the likelihood of a variant being the driver can be prioritized [23, 24]. It is worth noting that here, the clonal evolution was not inferred from time series sequencing data (which are difficult to obtain in cancer genomics), but the inherent constraints among variant frequencies due to the infinite sites assumption, (e.g., no clone can carry two variants unless the frequencies of one variant is always greater than the other; for details see [16]). Finally, similar to the clonal sequencing in LTEE, single cell sequencing data offers complementary information to clonal reconstruction in cancer genomics [25], and algorithms became available to infer tumor heterogeneity from low coverage single cell sequencing data [26, 27].

In this paper, we formalize the problem of clonal reconstruction from time course genomic sequencing data in a maximum likelihood framework, and devise a series of heuristic algorithms to address it. We further extend the algorithms to incorporate clonal sequencing data, aiming at reconstructing additional clones that are not sequenced. We simulated the bacterial population in long-term evolution experiments, and use the simulated genomic data to test our algorithms. The results show that the heuristic algorithms could accurately reconstruct as many clones as reconstructed by the brute-force algorithm or even better on average, while improving significantly on speed. We also discuss the effect of varying the number of clones in the population and the number of time points. Finally, we test our algorithms on a real LTEE dataset [4] from an *E. coli* population. Our algorithms successfully reconstruct clonal haplotypes that are not characterized by clonal sequencing, and reveal the evolutionary dynamics of the clones during the LTEE.

## Methods

### Modeling clonal evolution of bacteria

We model an evolving bacterial population using the clonal theory [28, 29], similar to the one used in cancer genomics [30]. We assume that all bacterial cells in an evolving population are descendants of a single *founding clone*. During the course of the evolution experiment, bacterial cells accumulate novel *mutations* forming new *clones*. In this study, we focus only on single nucleotide variations (SNVs); but the other types of variations (e.g., indels, structural variations and copy number variations) can be modelled in the same way. We further assume that the occurrences of mutations follow the *infinite sites assumption*, i.e., a mutation occurs at a single locus at most once during the period of evolution experiment.

The ancestral relationships between the clones in the evolving population can be represented as a *directed tree T*, referred to as the *clonal tree* in which the root represents the founder clone, every other node represents a clone introduced by one or more novel mutations, and each edge represents the direct ancestral relationships between the clones (Fig 1B). Each edge is labeled by the mutation(s) that distinguishes the *child* from its *parent*. When more than one mutation occurs during the evolution from the parent to the child, they can be clustered together and considered as a single mutation group. As a result, the *haplotype* of a clone (i.e., the variants contained in the clone) is represented by the path from the root to the node representing the clone.

The frequency of each clone at each specific time point is represented as a matrix *C* = [*c*_*ij*_], referred to as the *clonal frequency matrix (CFM)*, in which *c*_*i,j*_ indicates the frequency of clone *j* at the time point *i*. Our model assumes that the mutation occurs spontaneously; as a result, at any given time, the likelihood of a candidate clone to acquire a new mutation hence spawn a new clone is proportional to the frequency of the clone in the population at the time. The clonal tree *T* and the CFM *C* together can be depicted in a *Muller plot* [31] (Fig 1C), which is commonly used to visualize the evolutionary dynamics in a population [32].

### Time course clonal reconstruction problem

In order to monitor the evolutionary process in a bacterial population, metagenome sequencing can be conducted at a series of *N* time points, from which a *variant allele frequencies (VAF)* for all variation sites are obtained at each specific time point and represented as a *VAF matrix* [16], *F* = [*f*_*ij*_], where *f*_*i,j*_ indicates the allele frequency of the variant *j* at the time point *i*. Notably, each variant is first introduced by a mutation (or multiple mutations) at the time point *t*_*j*_, generating a novel clone (denoted by the specific mutation *j*) from its parent. Apparently, *t*_*j*_ is defined as the earliest time point *t*, such that *f*_*t,j*_ *>* 0, and for ∀*i < t, f*_*ij*_ = 0.

Given a VAF matrix *F*, our goal is to reconstruct the haplotype of each clone (i.e, the novel variants it contains) arising during the evolution experiment, or equivalently, to infer a clonal tree containing all observed mutations. Based on the clonal evolution model, we formally define the *time course clonal reconstruction problem* using a maximum likelihood formulation: given the input of matrix *F* = [*f*_*i,j*_] where 1 ≤ *i, j* ≤ *N* over *N* mutations (or novel clones) *sorted* over *N* time points (i.e., each mutation occurring at a known distinct time point), we want to find a directed tree *T* * = (*pr*(*i*), *i*), *i* = 1, 2, …, *N* on *N* nodes (where *pr*(*i*) is the *only* parent node of node *i*) that maximizes the following likelihood function,

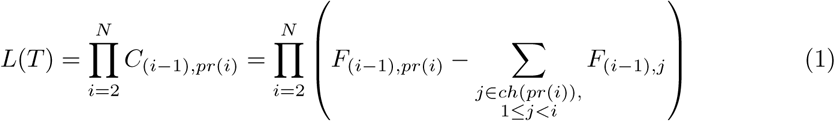

where *C*_*i,j*_ represents the (unknown) frequency of the clone *j* at the time point *i*, and *ch*(*i*) represents the set of all children of the node *i*. The likelihood function is computed by multiplying the likelihood of generating each clone in the clonal tree. As described above, the likelihood of generating the clone *i* or the probability of introducing the mutation *i* in clone *pr*(*i*) within the time segment between the points *i* − 1 and *i* is approximated by the frequency of the clone *pr*(*i*) at the time point *i* − 1, which can be computed as the frequency of the variant *pr*(*i*) subtracting the frequencies of all children of *pr*(*i*) born before the time point *i*.

We search for the optimal solution of a clonal tree *T* in the search space containing a total of (*N* − 1)! trees because at any given time *i* there are *i* − 1 putative parents to choose from. While some trees can be identified as invalid solutions when

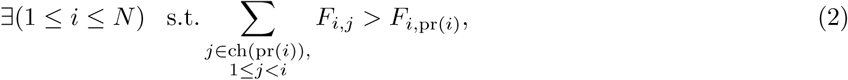

a brute force approach to search the ML solution in the entire tree space, referred to as the *exhaustive tree search algorithm* (ET; Algorithm 1) is still computationally expensive. Once the clonal tree is constructed, the haplotype of each clone (corresponding to a node in the tree) can be derived from the path from the root to the node. Note that a variant of this problem called the *variant allele frequency factorization problem (VAFFP)*, where the order (time) of appearance of each mutation is unknown and the likelihood assumption is not applicable, is proven to be NP-complete [16].

#### Algorithm 1 Exhaustive tree search algorithm (ET)

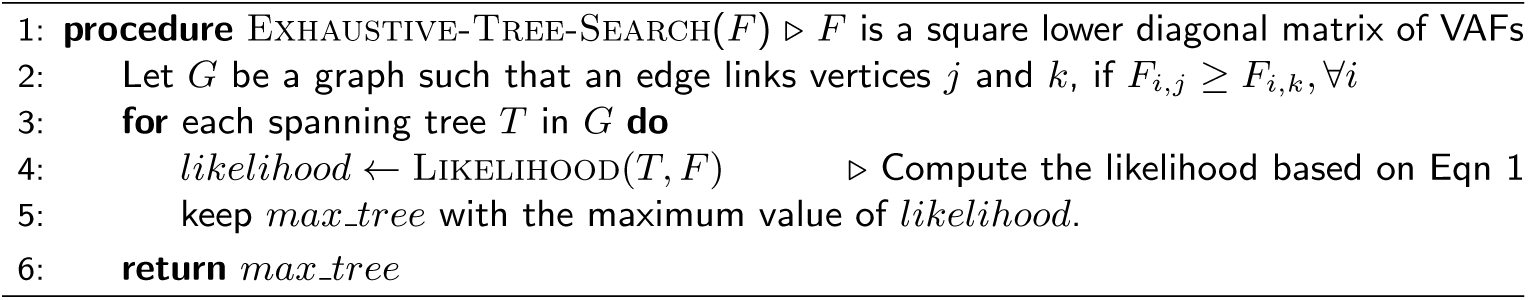

### Greedy tree search algorithm (GT)

To reduce the computational complexity of the *exhaustive tree search* (ET), we propose an algorithm using a greedy approach as follows (see Algorithm 2 for details). Start growing the directed tree from the root node (founder) such that at each iteration *i >* 1,

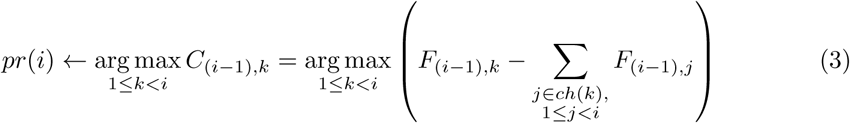

provided *pr*(*i*) does not lead to an invalid solution (according to equation 2). At any iteration *i*, if the assignment of *pr*(*i*) leads to an invalid solution, choose the next optimal choice at iteration *i* − 1, and continue the search. At any iteration *i*, if no more assignment leads to a valid solution, choose the next optimal choice at iteration *i* − 1 and continue the search until we find a valid *greedy solution* or until we run out of all choices (thus output no valid solution is found). Note that the worst case running time of this algorithm is still in *O*(*N* !) time, although in best case it runs in *O*(*N* ^2^) time.

#### Algorithm 2 Greedy tree search algorithm (GT)

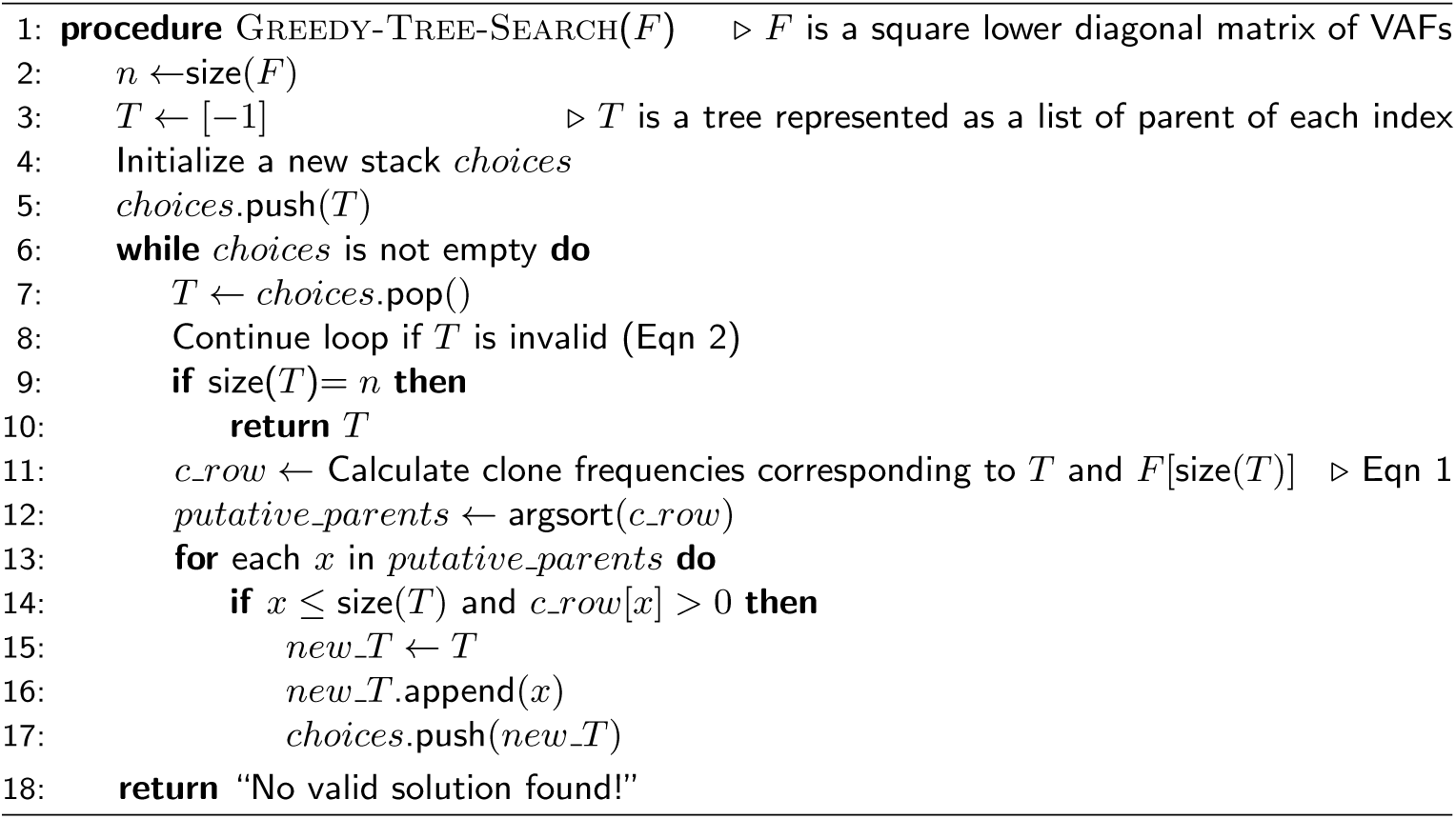

### Addressing sparse time course sequencing data

In practice, because of the often scattered genomic sequencing conducted in a time course, we may observe many mutations at the same time point. If the VAFs of some of these mutations are very similar across the time course, they are likely from the same clone, and thus can be grouped together and represented as a single mutation (group) as described above. If multiple mutations remain not grouped, but are all first observed at the same time point *t*, multiple clones should have emerged between the time points *t* − 1 and *t*. If the occurrence order of these mutations is determined, we may assume that the VAFs of all variants remain approximately constant between the time points *i* − 1 and *i*. Hence, we can simply extend the VAF matrix into a square lower diagonal matrix by introducing new rows between *t* − 1 and *t* for each mutation that is first observed at *t* while keeping the remaining VAFs constant. Then we can apply the greedy tree search (GT) algorithm to identify the ML clonal tree. In practice, as we do not know the occurrence order of these mutations, we have to also search for their optimal occurrence order among all possible permutations. An *exhaustive permutation search* (EP; Algorithm 3) would include 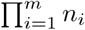! candidate permutations when there are *m* sets of unordered mutations with cardinalities *n*_1_, *n*_2_, …, *n*_*m*_, such that 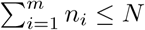. This is again computationally very expensive.

### Greedy permutation search (GP)

To reduce the computational complexity of the *exhaustive permutation search* (EP) algorithm, we propose a heuristic algorithm (for details see Algorithm 4) using a greedy approach as follows. For each set of unordered mutations *group*_*t*_ that first occur at the same time point *t*, extend the *F* matrix into a square lower diagonal matrix (up to time *t*) using each permutation of *group*_*t*_. Then, using the *greedy tree search* (GT; Algorithm 2) find the *ML tree* and the *ML score* for all permutations. The *ML permutation* at time *t* is determined as the one with the maximum ML score. At any iteration *i*, if the assignment of a permutation leads only to invalid trees, choose the next optimal choice at iteration *i* − 1, and continue the search until we find a valid tree. In the *greedy permutation search* algorithm, we only search 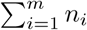! candidate matrices in the best case, but 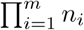! candidates in the worst case.

#### Algorithm 3 Exhaustive permutation search algorithm (EP)

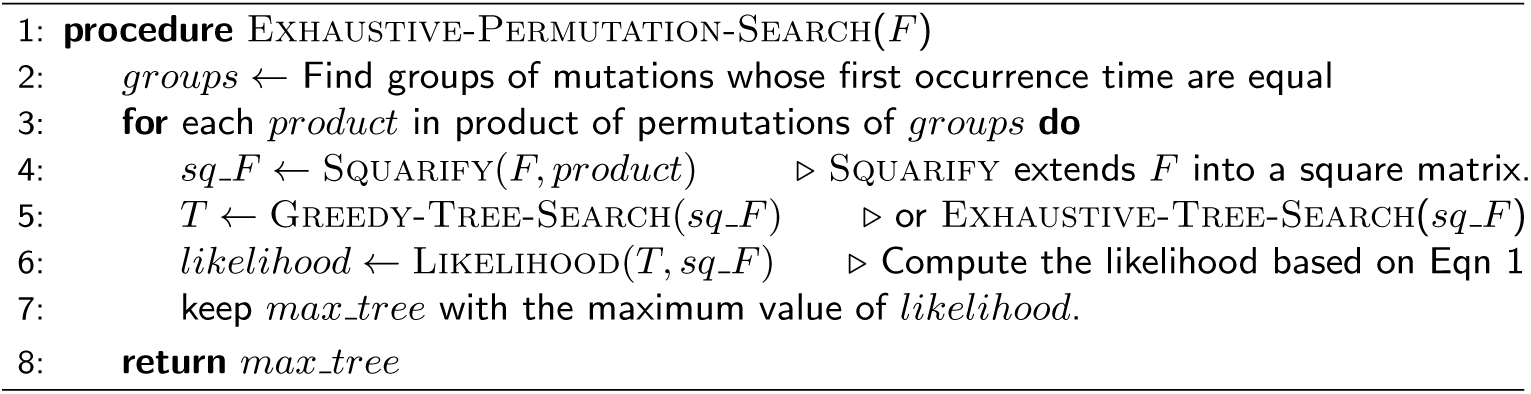

#### Algorithm 4 Greedy permutation search algorithm (GP)

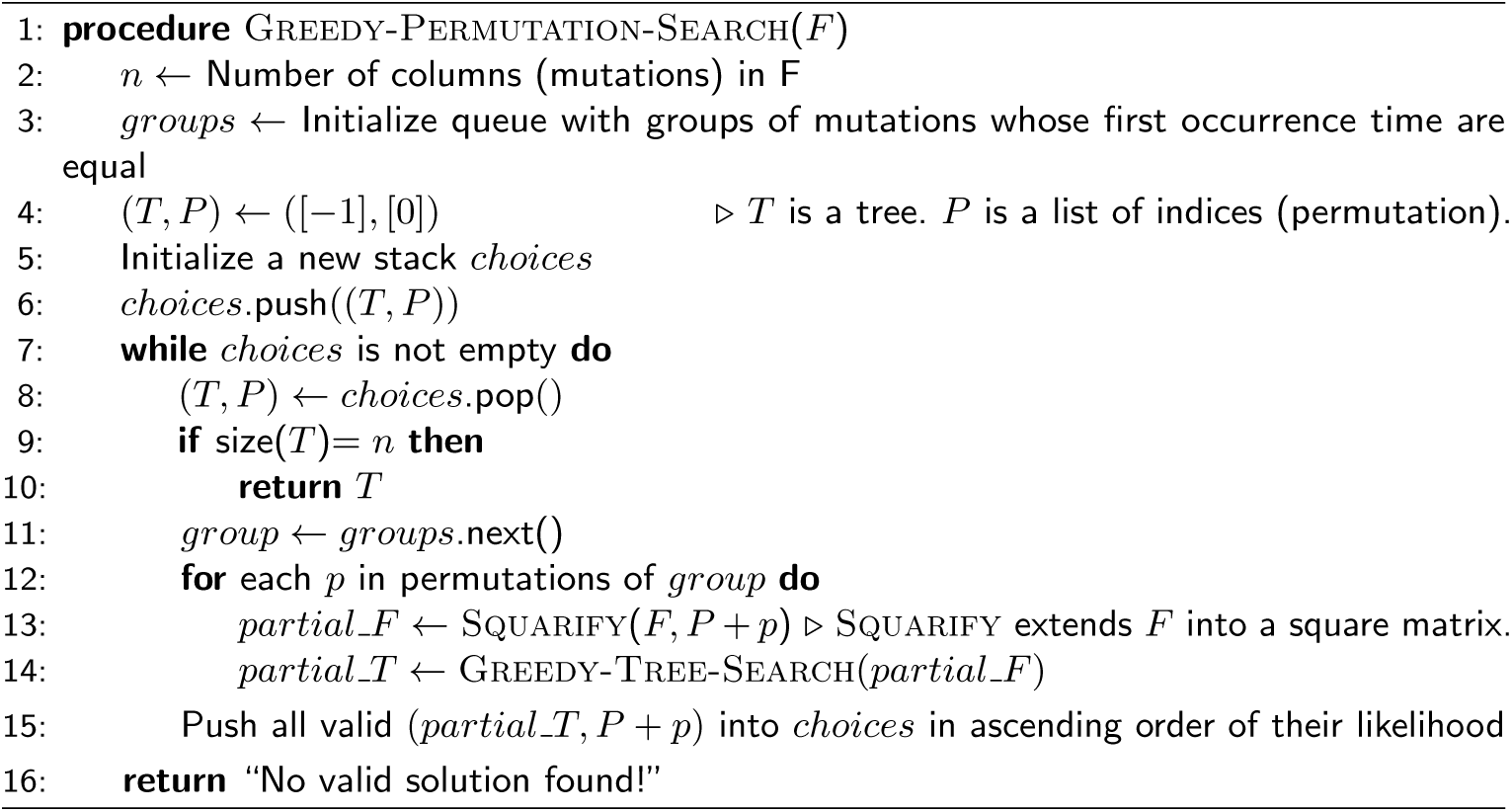

### Constrained search using sequenced clones

Sometimes we have additional information from the experiment when some randomly selected clones are sequenced during or at the end of the experiment. The haplotypes of these clones can be used to improve the search algorithm by enforcing that the clonal tree is consistent with the sequenced clones. There are two constraints that can be checked during the tree search to ensure the consistency. First, let the haplotypes of each sequenced clone (or the variants present in each clone) be represented as sets *A*_1_, *A*_2_, …*A*_*m*_. Then for each variant *v* in any sequenced clone, 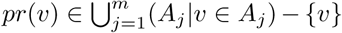. That is when we know the haplotype of at least one clone that contains variant *v*, its parent can only be one of the other variants in all the sequenced clones that has *v*. The second constraint is that for each pair of clones *A*_*i*_ and *A*_*j*_ and any variant *v* ∈ (*A*_*i*_ − *A*_*j*_) ∪ (*A*_*j*_ − *A*_*i*_), all variants *u* ∈ *A*_*i*_ ∩ *A*_*j*_ must appear before *v* in the path from the root node to *v* in the clonal tree. This constraint will make sure that the variants in the symmetric difference of two clones branch off after the common ancestral path is formed by the variants in the intersection of the two clones; otherwise, the infinite sites assumption will be violated.

### Metagenome sequencing data from an evolving *E. coli* population

We used data from the LTEE study on an *E. coli* population [4] to validate our methods. We used the paired-end Illumina sequencing reads data for the population 125, on which the metagenome sequencing was performed at six months interval during the course of three years, and eight clones were isolated and sequenced separately at the end of the experiment. We used Trimmomatic version 0.33 [33] to remove adapters and low quality bases and then mapped the reads to *E. coli* K12 MG1655 reference sequence (NC 000913.3) [34] with bwa-mem version 0.7.12 [35]. We removed reads supporting bases with forward/reverse read balance less than 0.25. Then we called variant sites where all of the following conditions satisfied: the VAF (approximated by the ratio of the number of reads supporting the variant allele to the sum of number of reads supporting the reference and variant alleles) was above 0.05, the sum of the number of reads supporting the variant and reference allele was above 10 and the number of variant reads was above 6. Then we removed inconsistent sites comparing the calls from different time points when the VAF at a site becomes zero and then non-zero again at a later time point. The VAFs were input to our algorithms to predict the clonal tree and the clonal frequencies, which were then visualized using Muller plot created using the R library ggmuller [36].

## Results

We compare the prediction performance of combinations of the two tree search algorithms (*ET* and *GT*) and the two permutation search algorithms (*EP* and *GP*), with two algorithms designed for tumor clonal reconstruction, AncesTree [16] and CITUP [21] on simulated data. *EP-ET* is the slowest algorithm. Hence, this algorithm is applied only to the case where the number of clones is very small. The other two combinations compared are *EP-GT* and *GP-GT*. The combination *GP-ET* does not differ much in performance with *GP-GT*. Therefore, we do not include the results from this combination here. We also compare the results with a baseline (or random) algorithm (*RP-RT*) where at each time-point *t* the parent is chosen at random from all the clones that have appeared before *t*. For non-square *F* matrices, a random permutation is chosen for each group of mutations that appear at the same time. The simulation procedure follows the clonal model described in methods. It starts with a founder clone and then at each new time point a new clone is introduced whose parent is chosen by random sampling from existing clones based on their frequencies in the population. The clonal frequencies are modified following a stochastic process between two consecutive time points. (Algorithm 5).

### Effect of number of clones

We generated 100 simulations for different number of clones − 10, 15, 20, 25, using the simulation algorithm. The number of time points at which the VAFs were observed is sampled from binomial distribution *B*(*n*, 0.6) where *n* is the number of clones. Fig 2 shows the distribution of recall (the proportion of clones correctly reconstructed by the algorithm), the running time in log scale and the log likelihood ratio of the predicted likelihood score over the true likelihood score. Since AncesTree and CITUP can output more than one solution, we calculated recall for each solution and used the maximum recall for comparison. The *GP-GT* algorithm correctly reconstructs almost as many or even more clones on average than *EP-ET* and *EP-GT* algorithms, while having a considerable (2-3 magnitudes) speed advantage (Fig 2A and 2B). The likelihood score returned by the *EP-ET* algorithm is the upper bound of any ML algorithm because this algorithm traverses the entire space of valid solutions and returns the tree that gives the maximum likelihood score. But the real clonal tree (following the simulation) may not be the same as the ML tree (Fig 2A and 2C). As the number of clones increases the likelihood scores returned by *EP-GT* deviates much further from the likelihood score of the true tree compared to the deviation of *GP-GT*, implying that the greedy heuristic not only helps in improving the speed but also reduces error in clonal reconstruction as the number of sequenced clones increases. Notably, AncesTree and CITUP do not take into consideration the sequential order of pooled sequencing data, and thus none of their reported trees are similar to the real clonal trees.

**Figure 2.**
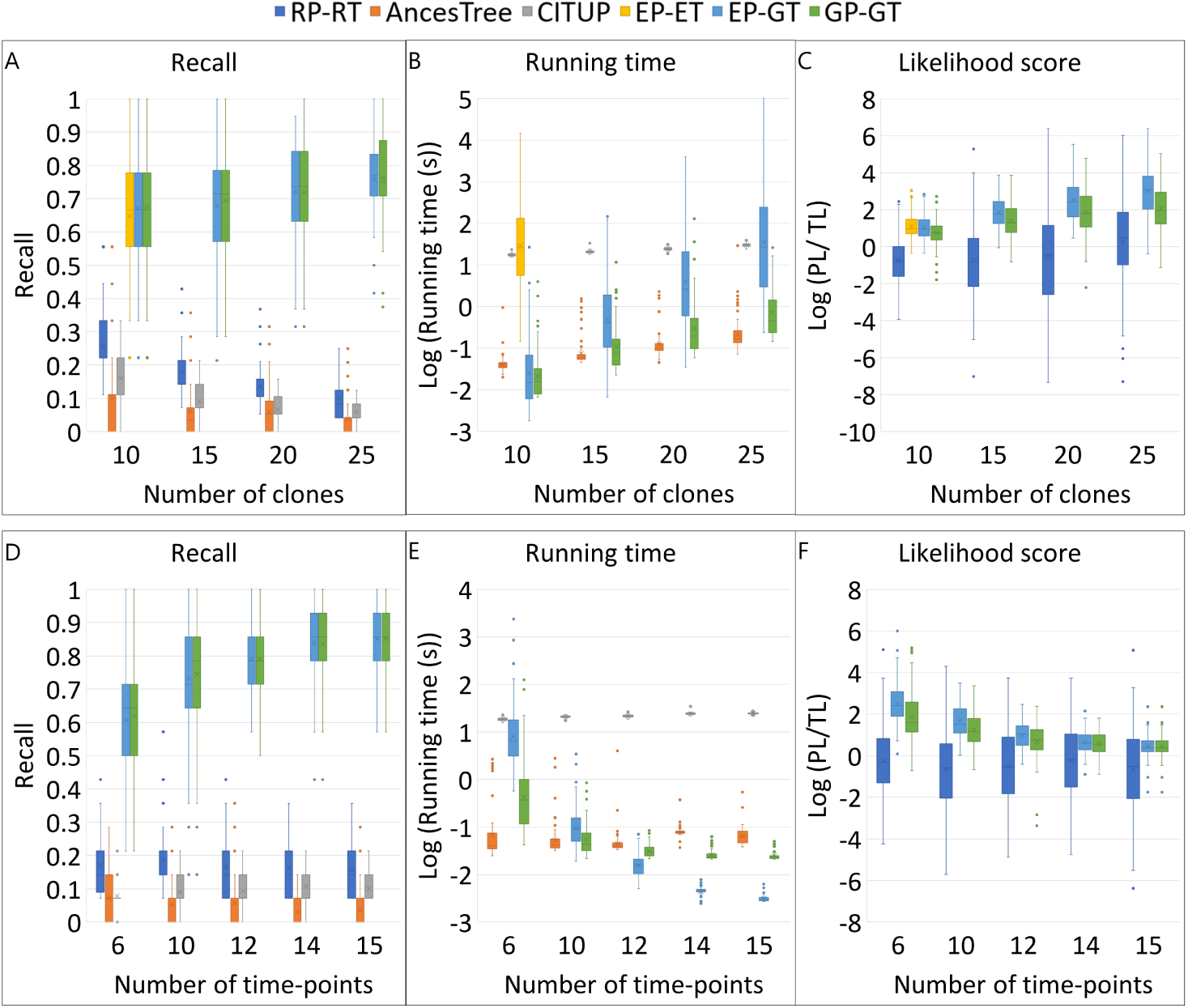
Comparison of recall, running time and likelihood scores of algorithms: RP-RT (Random Permutation search + Random Tree search), AncesTree, CITUP, EP-ET (Exhaustive Permutation search + Exhaustive Tree search), EP-GT (Exhaustive Permutation search + Greedy Tree search), GP-GT (Greedy Permutation search + Greedy Tree search). PL represents the likelihood score of the predicted clonal tree, and TL stands for the likelihood score of the true tree. Positive *Log*(*P L/TL*) indicates the algorithm predicted a clonal tree with greater likelihood than the real tree.

#### Algorithm 5 Simulation algorithm

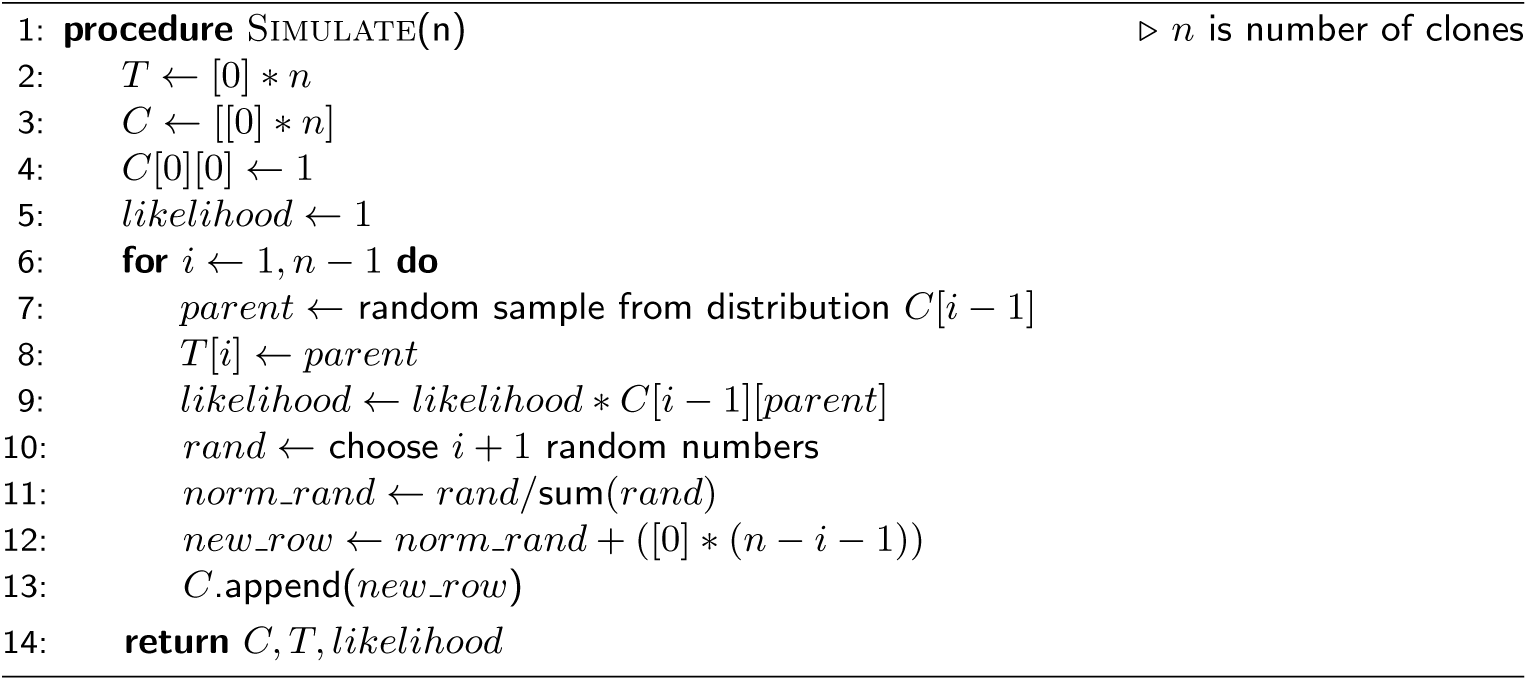

### Effect of sparse time course data

To study the effect of sparse time course sequencing data for a fixed number of clones, we generated 100 simulations with fixed number of clones (15), but varied the number of time points, where the VAFs were observed at 6, 10, 12, 14 and 15 time points, respectively. The results are shown in Figs 2D, 2E and 2F. As the number of time points increases, the number of unordered mutation groups reduces, which in turn reduces the size of the search space for the permutation search. As a result, the recall increases, reaching a maximum of about 0.85 on average when the number of time points is equal to the number of clones, for which no permutation search is needed.

### Constrained search

To test the effectiveness of constrained search given a set of sequenced clones we used the simulations generated earlier with 20 clones and compared the performance of the constrained search version of *GP-GT* algorithm by giving different number of true clones as input. We see that as the number of sequenced clones increases, the average number of misconstructed clones decreases, and so does the standard deviation (Fig 3).

**Figure 3.**
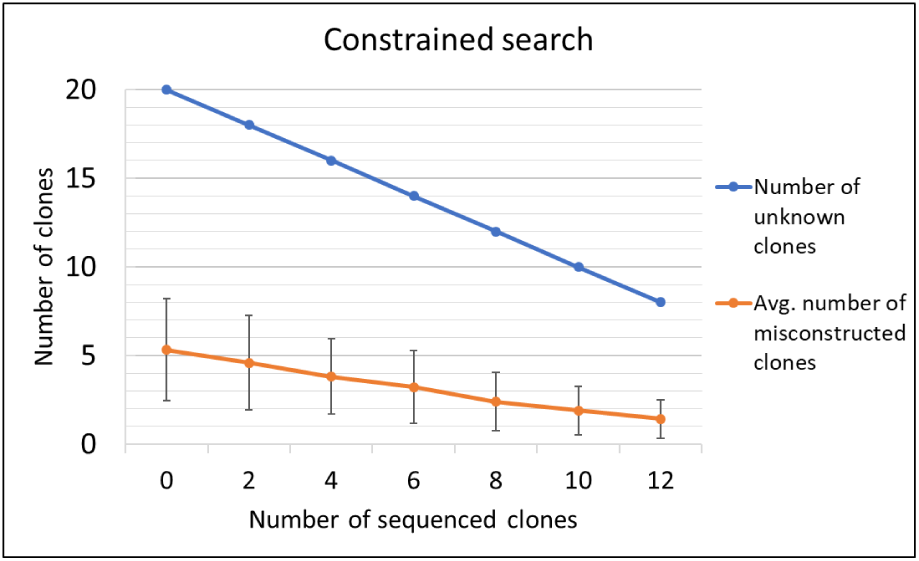
Results of constrained search on 100 simulations of 20 clones each.

### Analysis of metagenome sequencing data from the *E. coli* population

We used the metagenome sequencing data obtained from an LTEE study of an *E. coli* population [4], to validate our methods. It is to be noted that since the proportion of read support is used as an approximation for variant allele frequencies, these values are very noisy. When we did not allow any negative values in the clonal frequency matrix *C*, our algorithm did not return any valid solution. Thus, we relaxed this criterion to allow cells in *C* to have negative values greater than -0.4, which are then considered to have the frequency of 0. The input matrix *F* provided to the algorithm had VAF for 14 mutations at 6 time points. We use four haplotypes from eight sequenced clones (the other four being redundant clones with respect to the variants observed) in the constrained search using *GP-GT* [P]. The resulting clonal tree is shown in Fig 4A, which is consistent with sequenced clones (highlighted in gray). Fig 4B shows the haplotype of each clone and figure 4C shows the Muller plot showing the change in clonal frequencies over time. Note that the negative values in *C* were set to zero and then the rows were normalized to one. Fig 4D shows the clonal tree obtained when the known clones were not given as constraints. As shown in the simulation experiments, the accuracy of clonal reconstruction can be improved by including more time points, or by sequencing more clones.

**Figure 4.**
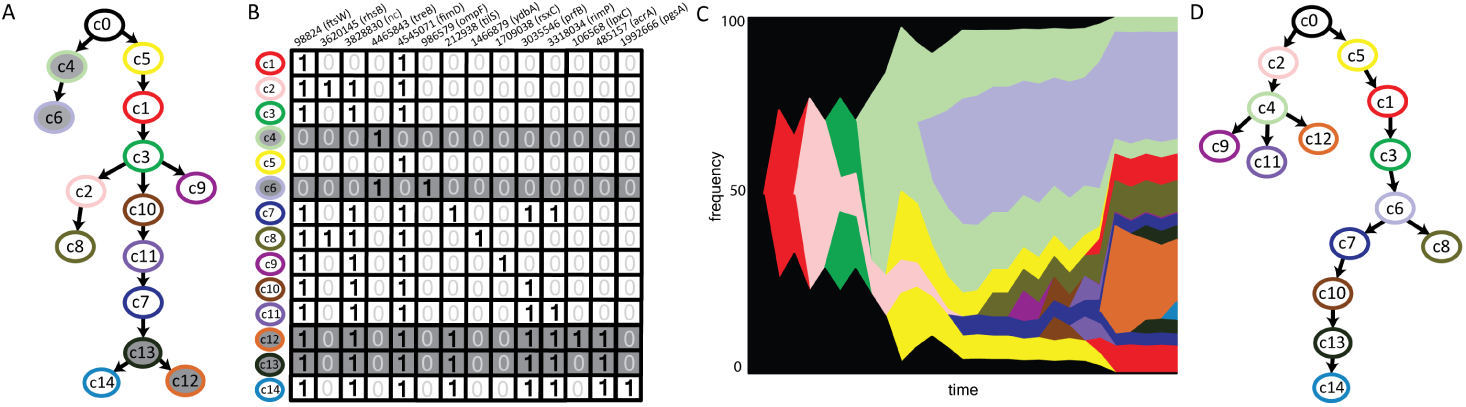
Clonal reconstruction on the *E. coli* LTEE data. (a) Clonal tree obtained by constrained search with *GP-GT* on the metagenome sequencing data. The four sequenced clones are shown in gray. (b) Reconstructed haplotypes depicted in a table where each row represents a clone and each column represents a variant. The variant locus and the gene where it is located are shown in the column header. Non coding region is marked as nc. When a clone has a specific variant, the corresponding cell is marked as 1, otherwise 0. Sequenced clones are highlighted in gray. (c) The Muller plot showing the predicted clonal frequency changes over time. The colors correspond to those in the clonal tree. (d) Clonal tree obtained without constrained search.

## Discussion

In this paper, we presented a maximum likelihood framework and a series of greedy-based heuristic algorithms to reconstruct the clonal haplotypes in a bacterial population from metagenome sequencing data obtained in a time course. All these algorithms can tolerate sparse time points sampling (thus multiple clones may arise in the same time period) while the constrained search algorithm can also incorporate clonal sequencing data as additional constraints for reconstructing un-sequenced clonal haplotypes. The results based on simulation experiment showed that, although the clones reconstructed by our algorithms are not identical to the real ones used in the simulation, they are highly similar, and more importantly, the likelihood computed on the reconstructed clones is comparable with (often higher than) the likelihood of the real ones, which implies that our algorithms achieved practically plausible optimal solutions under the maximum likelihood framework. Furthermore, our results demonstrated that the accuracy of clonal reconstruction can be improved by increasing the number of time points for metagenome sequencing or by increasing the number of sequenced clones. In particular, by sequencing more clones, not only the haplotypes of more clones can be directly derived, these derived haplotypes can impose additional constraints on the unknown (minor) haplotypes and thus improve the clonal reconstruction.

We note that the algorithms presented here report not only the haplotypes of clones, but also their frequencies over the time course. The next step after the clonal reconstruction is to identify the clones under selection during the course of evolution based on their frequencies. In this paper, we started to evaluate our algorithms on a relatively simple wild-type *E. coli* population. We plan to apply our algorithms to analyzing the time course genomic sequencing data from DNA repair deficient *E. coli* strains, in which hundreds of mutations occurred [4], to characterize the evolutionary dynamics of the complex population.

Although our algorithms were designed to analyze the sequencing data acquired from LTEE of bacterial populations (e.g., *E. coli*), it may have other applications such as in cancer genomics as described in the introduction. In addition, the metagenome sequencing approach was commonly adopted to study microbial communities containing hundreds of bacterial species, e.g., the human microbiome [37, 38] and the microbiome from natural habitats [39]. Recently, sequencing data acquired from the same microbial community at multiple time points become available [40, 41]. The current analyses of these data focus on the investigation of species and functional diversity in these communities. The computational approaches presented here can also be applied to these data, which will enable haplotype reconstruction of bacterial genomes and may reveal concerted evolution among bacterial species in the community. Interestingly, in the applications to both the cancer genomics and microbiome studies, clonal sequencing can be obtained through single cell sequencing, where our algorithm incorporating the clonal sequencing data can be directly applied.

## Conclusions

The main contribution of this paper is to develop a maximum likelihood framework to infer clonal evolutionary history from time course pooled sequencing data. The testing results on the simulation data show that our approach works better than the existing methods that do not take into consideration the sequential order of pooled sequencing data. The algorithms presented here is ready to be used for the analyses of sequencing data from large-scale LTEEs.

## List of abbreviations

LTEE: Long-term evolution experiment;
SNV: Single nucleotide variation;
WT: Wild-type;
CFM: Clonal frequency matrix;
VAF: Variant allele frequency;
ET: Exhaustive tree search;
GT: Greedy tree search;
RT: Random tree search;
EP: Exhausitve permutation search;
GP: Greedy permutation search;
RP: Random permutation search;
ML: Maximum likelihood.

## Acknowledgements

We thank Drs. Megan Behringer and Michael Lynch for very inspiring discussions.

## Declarations

### Availability of data and materials

The program (ClonalTREE) is available as open-source software on GitHub at https://github.com/COL-IU/ClonalTREE

### Ethics approval and consent to participate

Not applicable.

### Consent to publication

Not applicable.

### Author’s contributions

HT and WMI designed the study. WMI contributed tools for the analysis. WMI and HT analyzed the data, and wrote the paper. All authors read and approved the final manuscript.

## References

1. Elena, S.F., Lenski, R.E.: Microbial genetics: evolution experiments with microorganisms: the dynamics and genetic bases of adaptation. Nature Reviews Genetics 4(6), 457 (2003)

2. Rainey, P.B., Rainey, K.: Evolution of cooperation and conflict in experimental bacterial populations. Nature 425(6953), 72 (2003)

3. Barrick, J.E., Yu, D.S., Yoon, S.H., Jeong, H., Oh, T.K., Schneider, D., Lenski, R.E., Kim, J.F.: Genome evolution and adaptation in a long-term experiment with escherichia coli. Nature 461(7268), 1243 (2009)

4. Behringer, M.G., Choi, B.I., Miller, S.F., Doak, T.G., Karty, J.A., Guo, W., Lynch, M.: Escherichia coli cultures maintain stable subpopulation structure during long-term evolution. Proceedings of the National Academy of Sciences (2018)

5. Burke, M.K., Dunham, J.P., Shahrestani, P., Thornton, K.R., Rose, M.R., Long, A.D.: Genome-wide analysis of a long-term evolution experiment with drosophila. Nature 467(7315), 587 (2010)

6. Lenski, R.E., Rose, M.R., Simpson, S.C., Tadler, S.C.: Long-term experimental evolution in escherichia coli. i. adaptation and divergence during 2,000 generations. The American Naturalist 138(6), 1315–1341 (1991)

7. Sniegowski, P.D., Gerrish, P.J., Lenski, R.E.: Evolution of high mutation rates in experimental populations of e. coli. Nature 387(6634), 703 (1997)

8. Vasi, F., Travisano, M., Lenski, R.E.: Long-term experimental evolution in escherichia coli. ii. changes in life-history traits during adaptation to a seasonal environment. The american naturalist 144(3), 432–456 (1994)

9. Wielgoss, S., Barrick, J.E., Tenaillon, O., Cruveiller, S., Chane-Woon-Ming, B., Médigue, C., Lenski, R.E., Schneider, D.: Mutation rate inferred from synonymous substitutions in a long-term evolution experiment with escherichia coli. G3: Genes, Genomes, Genetics 1(3), 183–186 (2011)

10. Blount, Z.D., Barrick, J.E., Davidson, C.J., Lenski, R.E.: Genomic analysis of a key innovation in an experimental escherichia coli population. Nature 489(7417), 513 (2012)

11. Jewett, E.M., Steinrücken, M., Song, Y.S.: The effects of population size histories on estimates of selection coefficients from time-series genetic data. Molecular biology and evolution 33(11), 3002–3027 (2016)

12. Taus, T., Futschik, A., Schlötterer, C.: Quantifying selection with pool-seq time series data. Molecular biology and evolution 34(11), 3023–3034 (2017)

13. Perdigoto, C.: Cancer genomics: Tracking cancer evolution. Nature Reviews Genetics 18(7), 391 (2017)

14. McGranahan, N., Swanton, C.: Clonal heterogeneity and tumor evolution: past, present, and the future. Cell 168(4), 613–628 (2017)

15. Pon, J.R., Marra, M.A.: Driver and passenger mutations in cancer. Annual Review of Pathology: Mechanisms of Disease 10, 25–50 (2015)

16. El-Kebir M, A.-F.H.R.B. Oesper L: Reconstruction of clonal trees and tumor composition from multi-sample sequencing data. Bioinformatics 31(12) (2015)

17. Hajirasouliha, I., Mahmoody, A., Raphael, B.J.: A combinatorial approach for analyzing intra-tumor heterogeneity from high-throughput sequencing data. Bioinformatics 30(12), 78–86 (2014)

18. Deshwar, A.G., Vembu, S., Yung, C.K., Jang, G.H., Stein, L., Morris, Q.: Phylowgs: reconstructing subclonal composition and evolution from whole-genome sequencing of tumors. Genome biology 16(1), 35 (2015)

19. Donmez, N., Malikic, S., Wyatt, A.W., Gleave, M.E., Collins, C.C., Sahinalp, S.C.: Clonality inference from single tumor samples using low coverage sequence data. In: International Conference on Research in Computational Molecular Biology, pp. 83–94 (2016). Springer

20. McPherson, A.W., Roth, A., Ha, G., Chauve, C., Steif, A., de Souza, C.P., Eirew, P., Bouchard-Côté, A., Aparicio, S., Sahinalp, S.C., et al.: Remixt: clone-specific genomic structure estimation in cancer. Genome biology 18(1), 140 (2017)

21. McPherson, A.W., Sahinalp, C.S., Donmez, N., Malikic, S.: Clonality inference in multiple tumor samples using phylogeny. Bioinformatics 31(9), 1349–1356 (2015)

22. El-Kebir, M., Satas, G., Oesper, L., Raphael, B.J.: Inferring the mutational history of a tumor using multi-state perfect phylogeny mixtures. Cell systems 3(1), 43–53 (2016)

23. Qiao, Y., Quinlan, A.R., Jazaeri, A.A., Verhaak, R.G., Wheeler, D.A., Marth, G.T.: Subcloneseeker: a computational framework for reconstructing tumor clone structure for cancer variant interpretation and prioritization. Genome biology 15(8), 443 (2014)

24. Deveau, P., Colmet Daage, L., Oldridge, D., Bernard, V., Bellini, A., Chicard, M., Clement, N., Lapouble, E., Combaret, V., Boland, A., et al.: Quantumclone: clonal assessment of functional mutations in cancer based on a genotype-aware method for clonal reconstruction. Bioinformatics 34(11), 1808–1816 (2018)

25. Roerink, S.F., Sasaki, N., Lee-Six, H., Young, M.D., Alexandrov, L.B., Behjati, S., Mitchell, T.J., Grossmann, S., Lightfoot, H., Egan, D.A., et al.: Intra-tumour diversification in colorectal cancer at the single-cell level. Nature 556(7702), 457 (2018)

26. Navin, N.E.: Delineating cancer evolution with single cell sequencing. Science translational medicine 7(296), 296–29 (2015)

27. Ross, E.M., Markowetz, F.: Onconem: inferring tumor evolution from single-cell sequencing data. Genome biology 17(1), 69 (2016)

28. Tibayrenc, M., Kjellberg, F., Ayala, F.J.: A clonal theory of parasitic protozoa: the population structures of entamoeba, giardia, leishmania, naegleria, plasmodium, trichomonas, and trypanosoma and their medical and taxonomical consequences. Proceedings of the National Academy of Sciences 87(7), 2414–2418 (1990)

29. Shapiro, B.J.: How clonal are bacteria over time? Current opinion in microbiology 31, 116–123 (2016)

30. Nowell, P.C.: The clonal evolution of tumor cell populations. Science 194(4260), 23–28 (1976)

31. Muller, H.J.: Some genetic aspects of sex. The American Naturalist 66(703), 118–138 (1932)

32. Herron, M.D., Doebeli, M.: Parallel evolutionary dynamics of adaptive diversification in escherichia coli. PLoS biology 11(2), 1001490 (2013)

33. Bolger, A.M., Lohse, M., Usadel, B.: Trimmomatic: a flexible trimmer for illumina sequence data. Bioinformatics 30(15), 2114–2120 (2014)

34. Blattner, F.R., Plunkett, G., Bloch, C.A., Perna, N.T., Burland, V., Riley, M., Collado-Vides, J., Glasner, J.D., Rode, C.K., Mayhew, G.F., Gregor, J., Davis, N.W., Kirkpatrick, H.A., Goeden, M.A., Rose, D.J., Mau, B., Shao, Y.: The complete genome sequence of escherichia coli k-12. Science 277(5331), 1453–1462 (1997)

35. Li, H., Durbin, R.: Fast and accurate long-read alignment with burrows–wheeler transform. Bioinformatics 26(5), 589–595 (2010). doi:10.1093/bioinformatics/btp698

36. Noble, R.: R package: ggmuller. https://cran.r-project.org/package=ggmuller. Accessed: 2018-11-04

37. Huttenhower, C., Gevers, D., Knight, R., Abubucker, S., Badger, J.H., Chinwalla, A.T., Creasy, H.H., Earl, A.M., FitzGerald, M.G., Fulton, R.S., et al.: Structure, function and diversity of the healthy human microbiome. Nature 486(7402), 207 (2012)

38. Methé, B.A., Nelson, K.E., Pop, M., Creasy, H.H., Giglio, M.G., Huttenhower, C., Gevers, D., Petrosino, J.F., Abubucker, S., Badger, J.H., et al.: A framework for human microbiome research. Nature 486(7402), 215 (2012)

39. Gilbert, J.A., Jansson, J.K., Knight, R.: The earth microbiome project: successes and aspirations. BMC biology 12(1), 69 (2014)

40. Narayanasamy, S., Muller, E.E., Sheik, A.R., Wilmes, P.: Integrated omics for the identification of key functionalities in biological wastewater treatment microbial communities. Microbial biotechnology 8(3), 363–368 (2015)

41. Halfvarson, J., Brislawn, C.J., Lamendella, R., Vázquez-Baeza, Y., Walters, W.A., Bramer, L.M., D’Amato, M., Bonfiglio, F., McDonald, D., Gonzalez, A., et al.: Dynamics of the human gut microbiome in inflammatory bowel disease. Nature microbiology 2(5), 17004 (2017)

